# A stoichiometric selection advantage of DNA genomes over RNA genomes

**DOI:** 10.1101/2025.10.22.683889

**Authors:** Rosemary Yu

## Abstract

The RNA world hypothesis posits that RNA genomes preceded the universal use of DNA genomes in modern life. DNA genomes are commonly thought to have replaced RNA genomes due to greater chemical stability, however this is a misconception. This study investigates the selection pressures acting on the macromolecular stoichiometries of ribosome-containing cells (*ribocells*) during the epoch of the last universal common ancestor (LUCA), when DNA genomes and RNA genomes were in competition. Two selection pressures, (SP1) favoring stoichiometric simplicity of ribocell macromolecules (DNA, if any; RNA; and proteins), and (SP2) favoring minimal accumulation of unused/useless macromolecules in growing ribocells, were modeled as nonlinear optimization problems. Based on the current best estimates of the nature of LUCA, simulations showed that DNA genomes provided a robust selection advantage under these stoichiometric SPs. Moreover, results suggest that RNA-ssDNA hybrid genomes provided a stoichiometric selection advantage during the early LUCA epoch, consistent with a stepwise transition from RNA to DNA genomes. These findings indicate that DNA genomes were selected not for their inherent stability, but for their stoichiometric efficiency, resolving a long-standing question in the origin of life.

## Introduction

The RNA world hypothesis posits that RNA genomes, perhaps involving ssDNA as an intermediate (hybrid genomes) ^1,2^, preceded double-stranded DNA genomes used by all life on Earth today ^3-6^ (Fig 1A-C). There is a wide-spread notion that DNA was favoured over RNA as the genome storage material because it is more chemically stable (e.g. ^2,7^). This notion draws from the stability of the phosphodiester bond in DNA (half-life of 10^5^ to 10^7^ years ^8,9^), compared to RNA (half-life of 130 years ^10^). However, the N-glycosyl bond, which connects nucleobases to the sugar-phosphate backbone, has a half-life of only 100-200 years in DNA ^11^, while in RNA this bond has a half-life of about 6000 years ^12^. In other words, in a timeframe of tens to hundreds of years, the stability of DNA genomes and RNA genomes are actually comparable; what differs is only the type of degradation that predominates (fragmentation for RNA genomes, loss of nucleobases for DNA genomes). Therefore, the selective advantage of DNA genomes could not have been based on a higher genome stability, as has been noted before ^7,13^. Similarly, the apparent replication fidelity of DNA ^3,14,15^, low propensity to form secondary structures ^16^, and stability of the DNA double helix ^17,18^, have all been ruled out as the driving selection advantage for DNA genomes ^3,16^. To date, why DNA was selected over RNA as the genomic material – either once or multiple times independently ^2-4,15^ – remains unclear.

**Figure 1.**
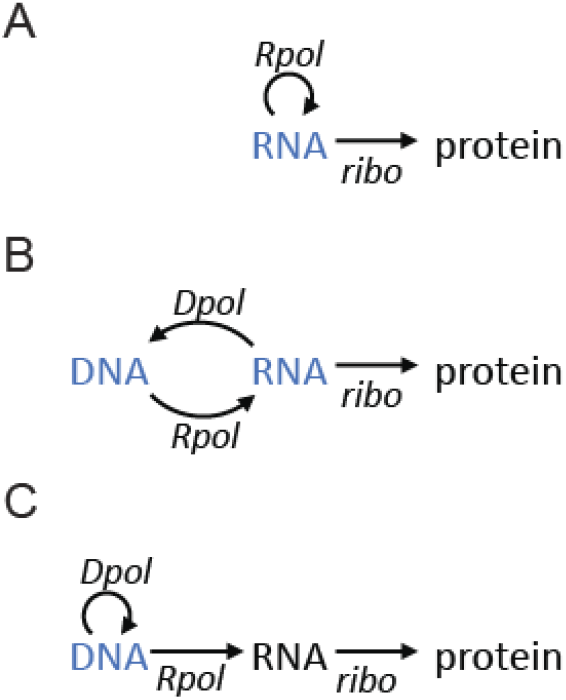
Three scenarios of genome materials in ribocells of the LUCA epoch. (A) RNA genome. (B) RNA-ssDNA hybrid genome. (C) dsDNA genome. Genome materials are indicated in blue. Rpol, RNA polymerase (either RNA-dependent or DNA-dependent); Dpol, DNA polymerase (either RNA-dependent or DNA-dependent); ribo, ribosomes.

The competition between DNA genomes and RNA genomes occurred in the epoch of the last universal common ancestor (LUCA), in a world of *ribocells* ^3^ that shared a common gene pool through pervasive lateral gene transfers. These ribocells were capable of transcription and translation ^2,3^, but likely divided in an uncontrolled manner through erratic membrane disturbances such as blebbing ^19,20^, with intracellular components segregating at random ^21^. This places two selection pressures on ribocells competing to successfully divide and disseminate. The first selection pressure (SP1) is for ribocells to be compositionally simple, i.e. requiring fewer total macromolecular components (DNA if any, RNA, proteins) for self-replication. This makes a new ribocell more likely to spontaneously assemble in a new vesicle with the right component stoichiometry ^22^. The second selection pressure (SP2) is for the ribocell to maintain the stoichiometry of its components while it grows (within the same vesicle) ^21^, to minimize the accumulation of unused/useless components. SP2 is equivalent to the selection for a more efficient use of environmental and intracellular resources, a constant pressure that continues to operate in single-celled organisms the modern day ^23,24^.

In this paper, I formulated SP1 and SP2 as non-linear optimization problems (NLPs), for each of the three genome scenarios in Fig 1A-C. Based on current best estimates of the nature of LUCA, including the likely sizes of intracellular components, simulations showed that DNA genomes have a robust selection advantage over RNA genomes under these stoichiometric selection pressures. RNA-ssDNA hybrid genomes were also advantageous under certain conditions, consistent with a stepwise transition from RNA to RNA-ssDNA to dsDNA genomes.

### Model description

Broadly speaking, the replication of the main macromolecular components of a self-replicating ribocell – DNA (if any), RNA, and proteins – are dependent on the activities of DNA/RNA polymerases or the ribosome. Coarse-graining these dependencies, we have:

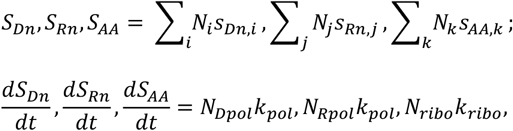

where *S*_*Dn*_,*S*_*Rn*_,*S*_*AA*_ represent the total number of DNA nucleotides, RNA nucleotides, and amino acids for all components in the ribocell; *N*_*i*_,*N*_*j*_,*N*_*k*_ represent the number (*N*) of each individual component (*i,j,k*) that are DNA, RNA, and proteins; *s*_*Dn,i*_,*s*_*Rn,j*_,*s*_*AA,k*_ represent the sizes (number of nucleotides or AAs) of each individual component (*i,j,k*); *N*_*Dpol*_,*N*_*Rpol*_,*N*_*ribo*_ represents the number (*N*) of DNA polymerases, RNA polymerases, and ribosomes; and *k*_*pol*_,*k*_*ribo*_ represents the polymerase and ribosome rate coefficients, in nucleotides per unit time and AAs per unit time, respectively. Then, the two stoichiometric selection pressures described above can be formulated as a non-linear optimization problem (NLP), in the form of:

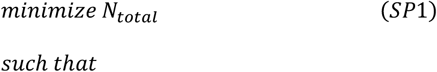

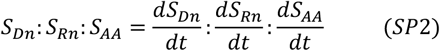

where *N*_*total*_ is the sum of all *N*_*i*_,*N*_*j*_,*N*_*k*_.

In the RNA genome scenario, ribosomes can interact productively (that is, the interaction can result in protein translation) with mRNA, and can also interact non-productively with the genome strand of RNA. In the NLP model for RNA genomes, 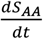 therefore reduces by a factor f:

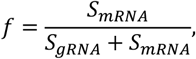

where *S* is the total number of nucleotides for each type of RNA as indicated. Note that the genome strand of RNA contains the complementary sequences of all genes, including not only mRNAs but also rRNA, tRNA, and/or ribozymes; this is elaborated in the Results section below. Because of this, *f* is always less than 0.5; the exact value of *f* varies depending on the (coarse-grained) sizes of each component. Similarly, in the DNA genome scenario, the RNA polymerase can interact productively with the complementary strand of DNA to produce mRNA, and can also interact non-productively with the coding strand. In the NLP model for DNA genomes, 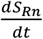 reduces by exactly 0.5, because the two strands of DNA are equal in size.

A detailed description of the formulation of each NLP model is given in the Methods section. MATLAB code of the models can be found at https://github.com/Radboud-YuLab/GeneticTakeover.

### The simplest ribocell – self-replicating polymerases and ribosomes

To run simulations on the NLP models for each genomic scenario (Fig 1A-C), it is necessary to consider the sizes of the components that ribocells should contain (number of nucleotides per DNA or RNA; number of AAs per protein). I began by considering the simplest self-replicating ribocell, containing only information processing and replication machineries: ribosomes and polymerases. The energy and chemical substrates needed for growth and dissemination were assumed to be readily available from the environment. Currently the ribosome of LUCA is thought to contain 4000-4500 nucleotides of rRNA sequence (in several fragments) ^25^, as well as 30-40 unique ribosomal proteins (RPs) of 50-100 AAs ^3,26-28^. The size of polymerases was assumed to be roughly the size of the catalytic module of RNA-dependent RNA polymerases of modern-day RNA viruses, at 500-600 AAs ^29^. The catalytic efficiencies of polymerases and ribosomes, *k*_*pol*_ and *k*_*ribo*_, were fixed to 36 nucleotides per unit time and 12 AA per unit time, respectively ^30,31^. In hybrid genomes and DNA genomes, both polymerases were assumed to be of the same size and have the same catalytic efficiencies.

The simulated optimal stoichiometry of polymerases and ribosomes for each genome scenario are given in Fig 2A and Table S1, showing that DNA genomes have a stoichiometric selection advantage over RNA genomes and hybrid genomes in 16% of occurrences. To account for the fact that, with a DNA genome, one molecule of (double-stranded) DNA contains twice as many nucleotides as one molecule of (single-stranded) genomic DNA or RNA in the other two scenarios, Fig 2B shows the molecular volumes for each scenario. As a frame of reference, the volume of unilamellar liposomes can be as small as 10^−6^ μm^332^; liposomes in the 10^−5^ to 10^−3^ μm^3^ range have been shown to readily encapsulate proteins and DNA ^33^, and replicate DNA by polymerase chain reaction ^34^. Therefore, simple self-replicating ribocells in the 10^−5^ μm^3^ (molecular volume; Fig 2B) to 10^−4^ μm^3^ (with water) range is a reasonable picture, though perhaps not a realistic one due to the assumption that energy and chemical substrates must be readily available without cellular metabolism – this is addressed in the following sections.

**Figure 2.**
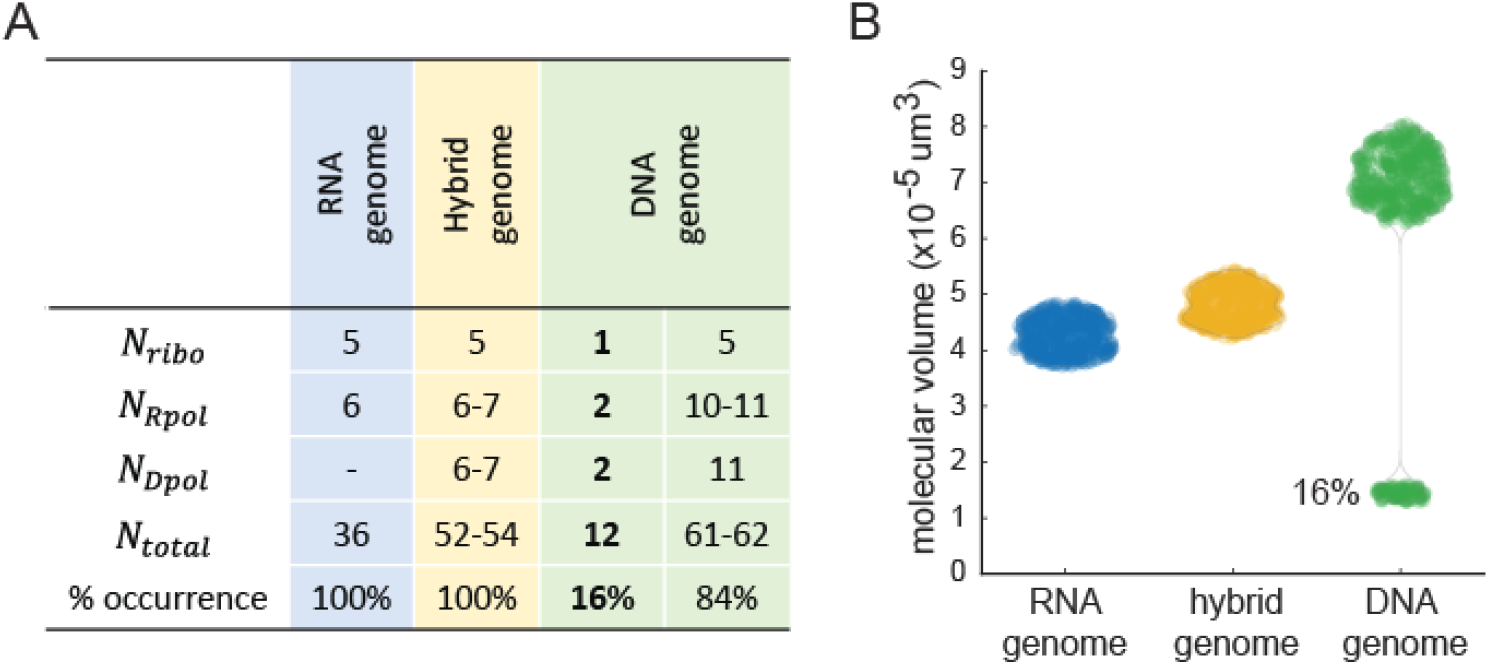
Stoichiometric selection advantage of DNA genomes in the simplest ribocell, containing only polymerases and ribosomes. (A) Model-simulated number of ribosomes, RNA polymerases, DNA polymerases, and total macromolecules (DNA if any, RNA, and protein) of ribocells in each genome scenario. The % occurrence of ribocells within each scenario is given. (B) Molecular volumes of ribocells with macromolecular stoichiometries as given in panel A.

A slightly more complex ribocell would contain not only polymerases and ribosomes, but also tRNAs and aminoacyl-tRNA synthetases (AARSes) to support ribosomal activities. Previous studies suggest that LUCA contained 44-45 tRNAs ^35^ and 17-20 AARSes ^36^. In terms of size, tRNA was assumed to be approximately the same size as modern day mature tRNA, at 70-90 nucleotides ^37^; AARSes were assumed to be the minimum size possible that retains catalytic activity, at around 100 AAs ^36^. With these additional components, simulations showed that ribocells with DNA genomes have a weak stoichiometric selection advantage over RNA genomes and hybrid genomes, at 7% of occurrences (Fig 3A and Table S2).

**Figure 3.**
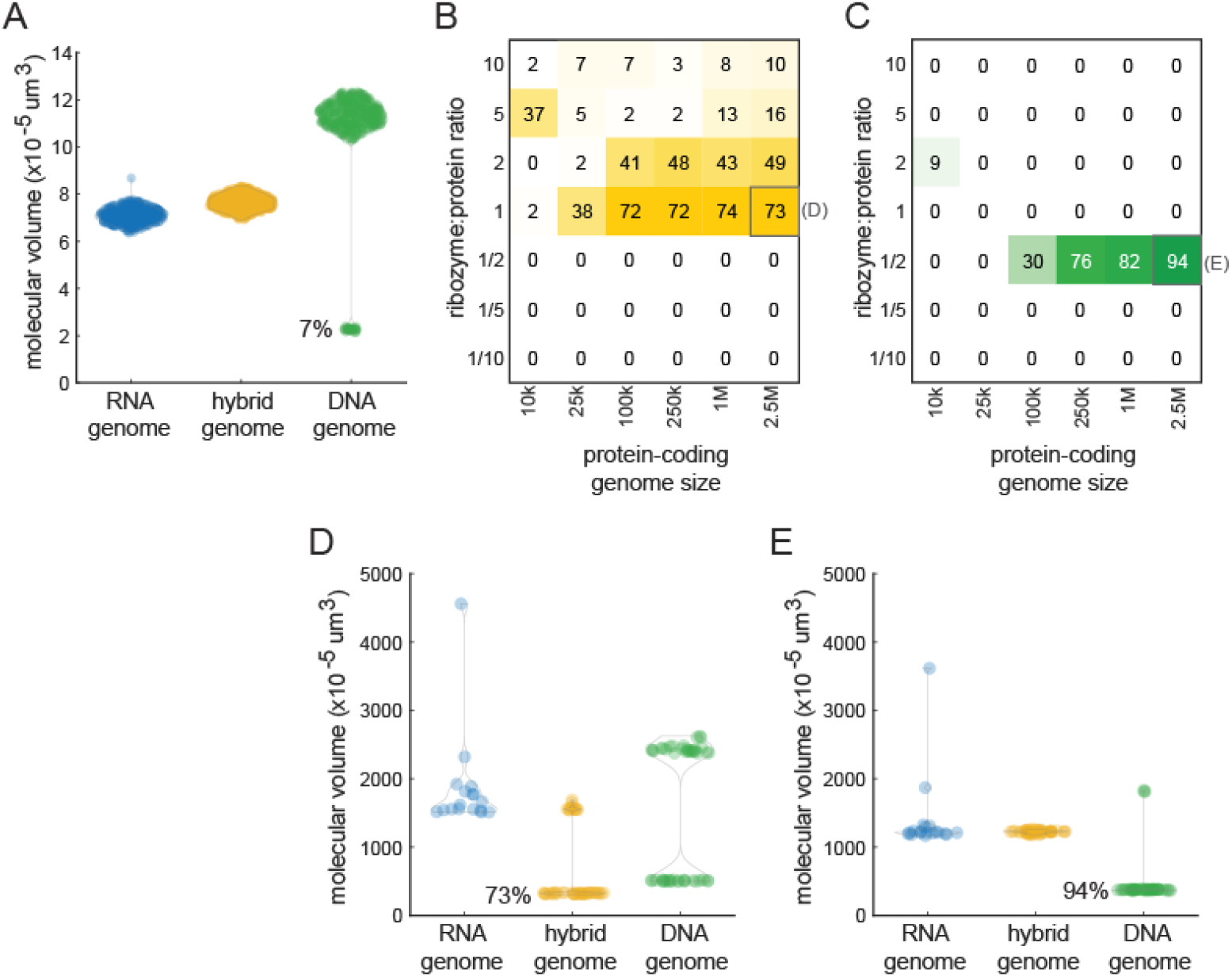
Stoichiometric selection advantage of hybrid genomes and DNA genomes in complex ribocells. (A) Molecular volumes of ribocells containing, in addition to ribosomes and polymerases, also AARSes and tRNAs. (B) The stoichiometric advantage of hybrid genomes with additional protein (enzyme) and ribozyme components. (C) The stoichiometric advantage of dsDNA genomes with additional protein (enzyme) and ribozyme components. (D) Molecular volumes of ribocells corresponding to the grey box in panel B. (E) Molecular volumes of ribocells corresponding to the grey box in panel C. Note that for all simulations so far, *k*_*pol*_ */ k*_*ribo*_ =3.

### Complex ribocells with additional protein and RNA components

To lift the assumption that all energy and chemical substrates needed for ribocell replication are readily available in the environment, I considered ribocells containing additional components that can carry out metabolic (or any other) functions. Of note, it is not necessary to specify each metabolic catalyst or reaction/pathway in the NLP models; an estimate of the coarse-grained sizes of catalysts that are functional proteins (enzymes) or RNA (ribozymes) is sufficient to run simulations. Recently the genome size of LUCA has been estimated to be 2.5-3 Mb, encoding around 2600 proteins ^38^, which is dramatically larger than earlier estimates of one to a few hundred proteins ^39-41^. An estimate of the abundance or diversity of ribozymes is not available, however it is reasonable to assume that evolution in the LUCA epoch have generally lead to (1) an increase in macromolecular diversity, accompanied by increased genome size, and (2) a transition from ribozymes (remnants of the RNA world) to enzymes and proteins as the main functional output of the genome ^41-43^.

I therefore scanned a range of protein-coding genome sizes up to 2.5Mb ^38^, as well as a range of ribozyme-to-protein ratios from 10 (more ribozymes than proteins) to 1/10 (more proteins than ribozymes). Fig 3B-C show that, as the genome size expands (left to right) and as protein-based enzymes replace RNA-based ribozymes (top to bottom), there is first a gradually increasing selection advantage for hybrid genomes (up to 74% of occurrences, Fig 3B and Table S3); then, as protein-based enzymes begin to outnumber RNA-based ribozymes, there is a strong stoichiometric selection advantage for DNA genomes (up to 94% of occurrences; Fig 3C and Table S3). The volumes of ribocells with these parameters are in the 10^−3^ μm^3^ (molecular volume; Fig 3D-E) to 10^−2^ μm^3^ (with water) range, comparable to modern day mycoplasma ^44^ and other ultra-small bacteria ^45^.

### Polymerase and ribosome efficiencies

The formulation of SP2 in the NLP models (see the Model Description section above) means that the exact values of *k*_*pol*_ and *k*_*ribo*_ have no effect on the simulation outcomes, but their relative value *k*_*pol*_ */ k*_*ribo*_ does. Thus far in the model simulations, I have fixed *k*_*pol*_ */ k*_*ribo*_ to 3, a ratio that is closely adhered to by modern day bacteria ^30^. In the LUCA epoch, this ratio is unknown and could undergo rapid changes, e.g. with random mutations in critical residues of polymerases and ribosomes. I therefore scanned a range of *k*_*pol*_ */ k*_*ribo*_ from 100 to 1/100. Fig 4A-C and Table S4 show that, if *k*_*pol*_ */ k*_*ribo*_, hybrid genomes (left panels, in yellow) have a moderate selection advantage (up to around 25% of occurrences). If *k*_*pol*_ */ k*_*ribo*_1 (Fig 4D-F and Table S4), DNA genomes (right panels, in green) have a strong selection advantage, particularly with larger genome sizes and more protein-coding genes; while hybrid genomes (left panels, in yellow) retain their moderate selection advantage at smaller genome sizes and more ribozyme-coding genes, as has been observed in Fig 3B-C.

**Figure 4.**
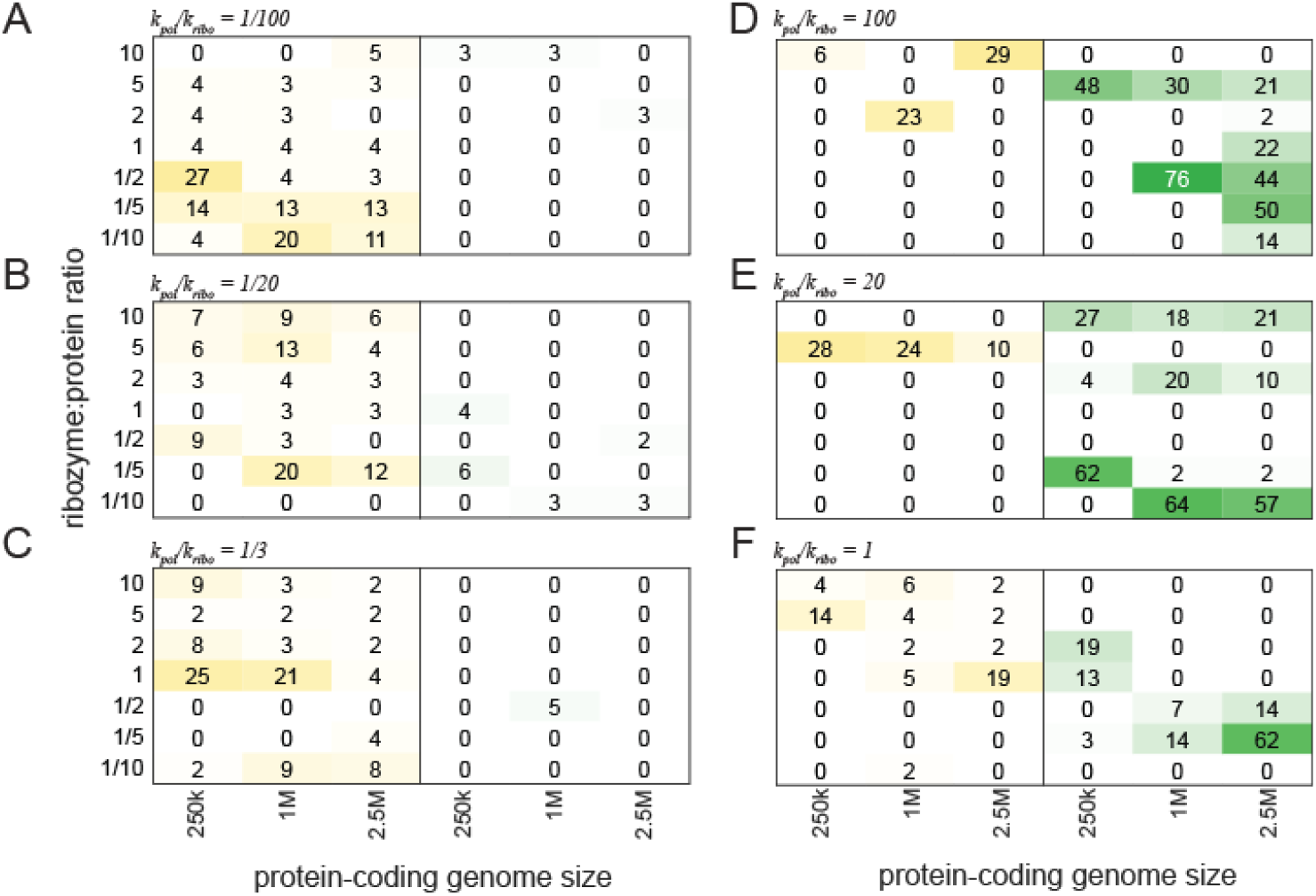
Stoichiometric selection advantage of hybrid genomes and DNA genomes in complex ribocells with different *k*_*pol*_ */ k*_*ribo*_ ratios. Each panel (A-F) corresponds to one *k*_*pol*_ */ k*_*ribo*_ ratio as indicated. Within each panel, the subpanel on the left (in yellow) represents stoichiometric advantage of hybrid genomes; the subpanel on the right (in green) represents stoichiometric advantage of DNA genomes. Note that *k*_*pol*_ */ k*_*ribo*_ =3.is already given in Figure 3B-C.

Taken together, these results indicate that the stoichiometric selection advantage of DNA genomes in the LUCA epoch is robust to changes in *k*_*pol*_ or *k*_*ribo*_, as long as (eventually) polymerases can synthesize DNA and RNA faster than ribosomes can synthesize proteins. Interestingly, these results also indicate that RNA-ssDNA hybrid genomes held a selective advantage in the early LUCA epoch, when (1) the genome sizes were still relatively small, (2) RNA-based ribozymes still outnumbered protein-based enzymes, and/or (3) *k*_*pol*_ */ k*_*ribo*_ have not yet stabilized to ≥ 1. This would be consistent with the idea that the transition from RNA genomes to DNA genomes may have taken place in two steps: first the ‘invention’ of DNA and its persistence in RNA-ssDNA hybrid genomes, for possibly a long period of time; followed by the ‘invention’ of DNA replication machineries, which may have occurred multiple times. This could explain why DNA genomes are universal in all extant life on Earth, while DNA replication, repair, and recombination machineries all appear to have arisen multiple times independently ^4,46,47^.

### Concluding remarks

The genomic material of LUCA remains a contentious topic. The results of this paper do not answer whether a DNA-based genome was selected for before or after LUCA, but rather resolve why it was selected for once it emerged. The common belief that DNA was favoured as the genetic material because it is more stable, is a misconception. This study shows that DNA genomes conferred a strong stoichiometric selection advantage to ribocells during the LUCA epoch, by promoting compositional simplicity (SP1) and reducing the accumulation of unused/useless macromolecular components, thereby allowing a more efficient use of intracellular and environmental resources (SP2).

## Methods

### NLP model 1. RNA genome

The variables in the RNA genome model (Fig 1A) are:

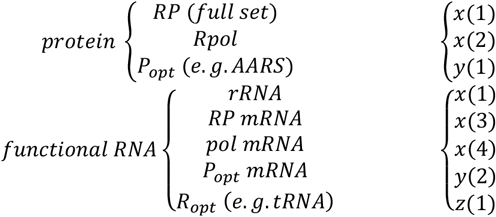

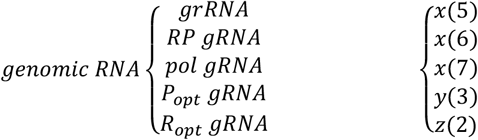

where *x* denotes the number of compulsory components of the ribocell (ribosome, RNA polymerase), *x* ≥ 1; *y* denotes the number of optional (opt) components with functional proteins (P), such as AARSes and enzymes, *y* = 0 if these optional components are not present, or *y* ≥ 1 if they are; and *z* denotes the number of optional components with functional RNA (R), such as tRNAs and ribozymes; *z* = 0 or *z* ≥ 1. Note that a functional ribosome consists of rRNA and a full set of RPs, thus these two components share the same variable, *x*(1).

The constraints of the RNA genome model are:

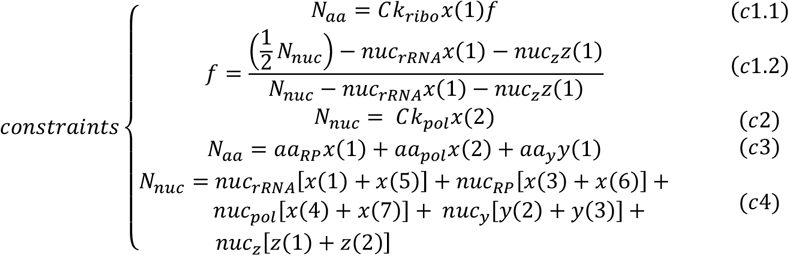

where *N*_*aa*_ and *N*_*nuc*_ denotes the total number of amino acids and nucleotides in the ribocell (in the Model Description section this is given by *S*);*C* is a positive number introduced to break up *SP*2 (see Model Description section) into equalities (*c*1 to *c*3, *c*2 to *c*4); and *f* stipulates that ribosomes will interact non-productively with all genomic RNA (including the genome strand of rRNA, tRNA, and ribozymes), but will not interact with the functional strand of rRNA, tRNA, and ribozymes, as described in the Model Description section. Note that here *f* is given explicitly for clarity and space; *c*1.1 and *c*1.2 are in fact two components of the same constraint *c*1. The number of amino acids of any protein component, *aa*_∗_, is 1/3 the number of nucleotides of the RNA of that component, *nuc*_∗_. The objective of the NLP (*SP*1, see Model Description) is to minimize the total number of molecules:

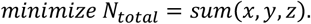

Since the NLP is underdetermined with respect to the number of functional RNA components and their corresponding genomic RNA components (e.g. *x*(3) and *x*(6)), an additional constraint is then applied,

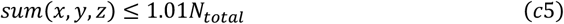

where the flexibilization factor 1.01 is introduced to prevent floating-point roundoff errors. A new objective is then set to minimize the number of genomic RNA molecules ^48^:

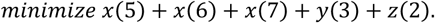

### NLP model 2. hybrid genome

The variables in the hybrid genome model (Fig 1B) are:

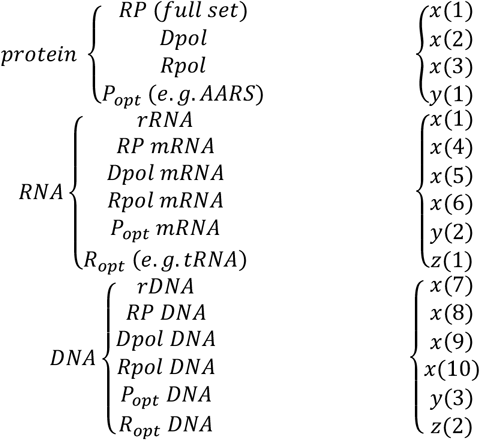

The difference here compared to the RNA genome is that there are now two polymerases, a DNA-directed RNA polymerase (*Rpol*) and a reverse transcriptase (*Dpol*), see Fig 1B.

The constraints of the hybrid genome model are:

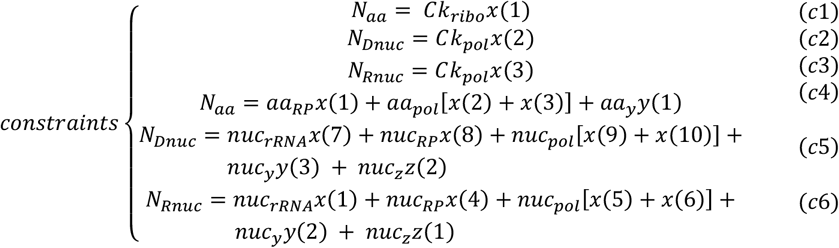

As with the RNA genome model, the first objective of the NLP is to minimize the total number of molecules:

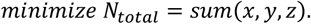

Then, an additional constraint is applied:

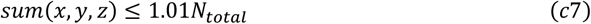

Although this is a hybrid genome, to address the issue that the NLP is underdetermined with respect to the number of RNA components and their corresponding genomic ssDNA components, here we consider the only ssDNA molecules to be the ‘genome strand’ to be minimized ^48^:

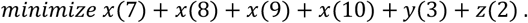

### NLP model 3. DNA genome

The variables in the DNA genome model (Fig 1C) are identical to the variables in the hybrid genome model; note however that *Dpol* in this model refers to the DNA polymerase (contrast Fig 1B and Fig 1C). The DNA molecules in the DNA genome is double-stranded, so the sizes of each molecule doubles, but the number of each molecule (variables *x,y,z*) are the same.

The constraints of the DNA genome model are:

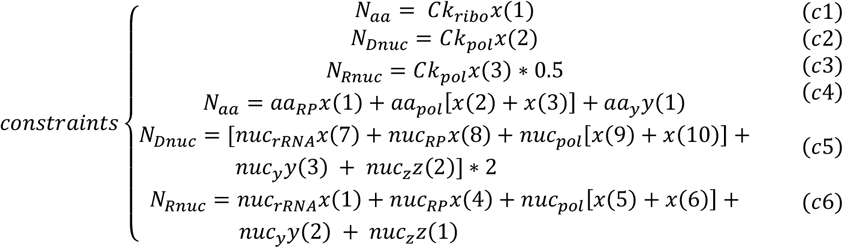

The double-stranded nature of the DNA genome is reflected in *c*5. Additionally as described in the ‘Model description’ section, in this scenario the RNA polymerase *x*(3) can interact with both strands of the DNA, but only half of this interaction is productive; therefore a factor of exactly 0.5 is introduced in *c*3.

The first objective of the NLP is to minimize the total number of molecules:

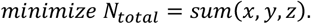

Then, an additional constraint is applied:

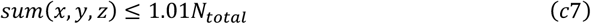

And a new objective is then set to minimize the number of DNA molecules ^48^:

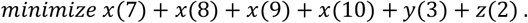

### Model simulations

The models for each scenario (Fig 1A-C) were simulated with the following input; see Results section for an explanation of the choice of the input values and their references. For the last two input values, recall that *y* denotes the number of optional (opt) components with functional proteins, such as AARSes and enzymes; and *z* denotes the number of optional components with functional RNA, such as tRNAs and ribozymes. Input 1a corresponds to the simplest ribocell with only polymerases and ribosomes; input 2 corresponds to the addition of AARSes and tRNAs; input 3 corresponds to the addition of proteins (enzymes) and ribozymes; input 4 corresponds to the evaluation of polymerase and ribosome efficiencies.

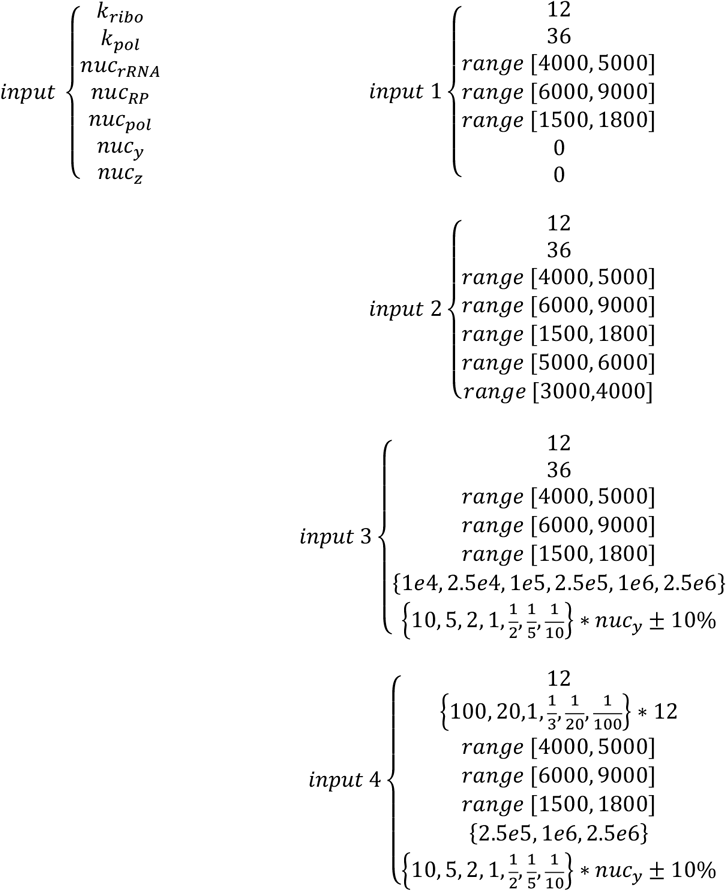

Recall that *nuc*_∗_ refers to the number of nucleotides of the RNA of that component, which is used to calculate *aa*_∗_ (where appropriate) by a factor of 1/3. Where a range is indicated, a random integer is selected within the range 50 to 500 times; where a flexibilization factor (±10%) is indicated, a random float is selected with the flexibilization range 50 to 500 times. If the round-to-integers flag is raised, first each value of the solution (variables *x,y,z*) are rounded to the nearest 0.2, then the smallest integer values for the full set of solution is sought while preserving the stoichiometry of the values.

For molecular volume calculations, an average molecular volume of 300Å^3^ was used for both DNA and RNA nucleotides ^49^; an average molecular volume of 140Å^3^ was used for AAs ^50^.

### Computational resources

All procedures are implemented in MATLAB R2025a with the Statistics and Machine Learning Toolbox; Bioinformatics Toolbox; Optimization Toolbox; Global Optimization Toolbox; Symbolic Math Toolbox; and Parallel Computing Toolbox. On a personal computer with a 10th Gen Intel(R) Core(TM) i7-10750H, 2.60 GHz, 6 cores, 12 logical processors, and 32 GB RAM, the processing time for the entire workflow is about 1.5 hours.

## Code availability

The NLP models are written in MATLAB and openly available at https://github.com/Radboud-YuLab/GeneticTakeover. The scripts used in this paper for model simulations, data analysis, and visualizations, as well as all supplemental tables and figures, are accessible at https://github.com/Radboud-YuLab/GeneticTakeover.

## Notes

https://github.com/Radboud-YuLab/GeneticTakeover

### Competing Interest Statement

The authors have declared no competing interest.

## References

1 Weiss, M. C., Preiner, M., Xavier, J. C., Zimorski, V. & Martin, W. F. The last universal common ancestor between ancient Earth chemistry and the onset of genetics. PLoS Genet 14, e1007518 (2018). 10.1371/journal.pgen.1007518

2 Koonin, E. V. & Martin, W. On the origin of genomes and cells within inorganic compartments. Trends Genet 21, 647–654 (2005). 10.1016/j.tig.2005.09.006

3 Forterre, P. The Last Universal Common Ancestor of Ribosome-Encoding Organisms: Portrait of LUCA. J Mol Evol 92, 550–583 (2024). 10.1007/s00239-024-10186-9

4 Leipe, D. D., Aravind, L. & Koonin, E. V. Did DNA replication evolve twice independently? Nucleic Acids Res 27, 3389–3401 (1999). 10.1093/nar/27.17.3389

5 Koonin, E. V., Krupovic, M., Ishino, S. & Ishino, Y. The replication machinery of LUCA: common origin of DNA replication and transcription. BMC Biol 18, 61 (2020). 10.1186/s12915-020-00800-9

6 Pereira Dos Santos Junior, A., Jose, M. V. & Torres de Farias, S. From RNA to DNA: Insights about the transition of informational molecule in the biological systems based on the structural proximity between the polymerases. Biosystems 206, 104442 (2021). 10.1016/j.biosystems.2021.104442

7 Forterre, P. The two ages of the RNA world, and the transition to the DNA world: a story of viruses and cells. Biochimie 87, 793–803 (2005). 10.1016/j.biochi.2005.03.015

8 Schroeder, G. K., Lad, C., Wyman, P., Williams, N. H. & Wolfenden, R. The time required for water attack at the phosphorus atom of simple phosphodiesters and of DNA. Proc Natl Acad Sci U S A 103, 4052–4055 (2006). 10.1073/pnas.0510879103

9 Matange, K., Tuck, J. M. & Keung, A. J. DNA stability: a central design consideration for DNA data storage systems. Nat Commun 12, 1358 (2021). 10.1038/s41467-021-21587-5

10 Li, Y. & Breaker, R. R. Kinetics of RNA Degradation by Specific Base Catalysis of Transesterification Involving the 2’-Hydroxyl Group. Journal of the American Chemical Society 121, 5364–5372 (1999). 10.1021/ja990592p

11 Schroeder, G. K. & Wolfenden, R. Rates of spontaneous disintegration of DNA and the rate enhancements produced by DNA glycosylases and deaminases. Biochemistry 46, 13638–13647 (2007). 10.1021/bi701480f

12 Stockbridge, R. B., Schroeder, G. K. & Wolfenden, R. The rate of spontaneous cleavage of the glycosidic bond of adenosine. Bioorg Chem 38, 224–228 (2010). 10.1016/j.bioorg.2010.05.003

13 Takeuchi, N., Hogeweg, P. & Koonin, E. V. On the origin of DNA genomes: evolution of the division of labor between template and catalyst in model replicator systems. PLoS Comput Biol 7, e1002024 (2011). 10.1371/journal.pcbi.1002024

14 Leu, K., Obermayer, B., Rajamani, S., Gerland, U. & Chen, I. A. The prebiotic evolutionary advantage of transferring genetic information from RNA to DNA. Nucleic Acids Res 39, 8135–8147 (2011). 10.1093/nar/gkr525

15 Poole, A. M. & Logan, D. T. Modern mRNA proofreading and repair: clues that the last universal common ancestor possessed an RNA genome? Mol Biol Evol 22, 1444–1455 (2005). 10.1093/molbev/msi132

16 Ma, W., Yu, C., Zhang, W., Wu, S. & Feng, Y. The emergence of DNA in the RNA world: an in silico simulation study of genetic takeover. BMC Evol Biol 15, 272 (2015). 10.1186/s12862-015-0548-1

17 Wienken, C. J., Baaske, P., Duhr, S. & Braun, D. Thermophoretic melting curves quantify the conformation and stability of RNA and DNA. Nucleic Acids Res 39, e52 (2011). 10.1093/nar/gkr035

18 Zhang, K., Hodge, J., Chatterjee, A., Moon, T. S. & Parker, K. M. Duplex Structure of Double-Stranded RNA Provides Stability against Hydrolysis Relative to Single-Stranded RNA. Environ Sci Technol 55, 8045–8053 (2021). 10.1021/acs.est.1c01255

19 Koonin, E. V. & Mulkidjanian, A. Y. Evolution of cell division: from shear mechanics to complex molecular machineries. Cell 152, 942–944 (2013). 10.1016/j.cell.2013.02.008

20 Briers, Y., Walde, P., Schuppler, M. & Loessner, M. J. How did bacterial ancestors reproduce? Lessons from L-form cells and giant lipid vesicles: multiplication similarities between lipid vesicles and L-form bacteria. Bioessays 34, 1078–1084 (2012). 10.1002/bies.201200080

21 Goldman, A. D. & Landweber, L. F. Oxytricha as a modern analog of ancient genome evolution. Trends Genet 28, 382–388 (2012). 10.1016/j.tig.2012.03.010

22 Monnard, P. A. & Deamer, D. W. Membrane self-assembly processes: steps toward the first cellular life. Anat Rec 268, 196–207 (2002). 10.1002/ar.10154

23 Yu, R. et al. Nitrogen limitation reveals large reserves in metabolic and translational capacities of yeast. Nat Commun 11, 1881 (2020). 10.1038/s41467-020-15749-0

24 Basan, M. et al. Overflow metabolism in Escherichia coli results from efficient proteome allocation. Nature 528, 99–104 (2015). 10.1038/nature15765

25 Men, Y., Lu, G., Wang, Y., Lin, J. & Xie, Q. Reconstruction of the rRNA Sequences of LUCA, with Bioinformatic Implication of the Local Similarities Shared by Them. Biology (Basel) 11 (2022). 10.3390/biology11060837

26 Korobeinikova, A. V., Garber, M. B. & Gongadze, G. M. Ribosomal proteins: structure, function, and evolution. Biochemistry (Mosc) 77, 562–574 (2012). 10.1134/S0006297912060028

27 Mushegian, A. Protein content of minimal and ancestral ribosome. RNA 11, 1400–1406 (2005). 10.1261/rna.2180205

28 Melnikov, S., Manakongtreecheep, K. & Soll, D. Revising the Structural Diversity of Ribosomal Proteins Across the Three Domains of Life. Mol Biol Evol 35, 1588–1598 (2018). 10.1093/molbev/msy021

29 Jia, H. & Gong, P. A Structure-Function Diversity Survey of the RNA-Dependent RNA Polymerases From the Positive-Strand RNA Viruses. Front Microbiol 10, 1945 (2019). 10.3389/fmicb.2019.01945

30 Proshkin, S., Rahmouni, A. R., Mironov, A. & Nudler, E. Cooperation between translating ribosomes and RNA polymerase in transcription elongation. Science 328, 504–508 (2010). 10.1126/science.1184939

31 Fitzsimmons, W. J. et al. A speed-fidelity trade-off determines the mutation rate and virulence of an RNA virus. PLoS Biol 16, e2006459 (2018). 10.1371/journal.pbio.2006459

32 Jesorka, A. & Orwar, O. Liposomes: technologies and analytical applications. Annu Rev Anal Chem (Palo Alto Calif) 1, 801–832 (2008). 10.1146/annurev.anchem.1.031207.112747

33 Burger, M., Brigger, F., Mantella, V. & Leroux, J. C. Encapsulation of protein/DNA complexes into unilamellar liposomes via annexin-mediated membrane recruitment and sonication. Cell Rep Methods 5, 101073 (2025). 10.1016/j.crmeth.2025.101073

34 Oberholzer, T., Albrizio, M. & Luisi, P. L. Polymerase chain reaction in liposomes. Chem Biol 2, 677–682 (1995). 10.1016/1074-5521(95)90031-4

35 van der Gulik, P. T. & Hoff, W. D. Anticodon Modifications in the tRNA Set of LUCA and the Fundamental Regularity in the Standard Genetic Code. PLoS One 11, e0158342 (2016). 10.1371/journal.pone.0158342

36 Rubio Gomez, M. A. & Ibba, M. Aminoacyl-tRNA synthetases. RNA 26, 910–936 (2020). 10.1261/rna.071720.119

37 Santos, F. B. & Del-Bem, L. E. The Evolution of tRNA Copy Number and Repertoire in Cellular Life. Genes (Basel) 14 (2022). 10.3390/genes14010027

38 Moody, E. R. R. et al. The nature of the last universal common ancestor and its impact on the early Earth system. Nat Ecol Evol 8, 1654–1666 (2024). 10.1038/s41559-024-02461-1

39 Weiss, M. C. et al. The physiology and habitat of the last universal common ancestor. Nat Microbiol 1, 16116 (2016). 10.1038/nmicrobiol.2016.116

40 Berkemer, S. J. & McGlynn, S. E. A New Analysis of Archaea-Bacteria Domain Separation: Variable Phylogenetic Distance and the Tempo of Early Evolution. Mol Biol Evol 37, 2332–2340 (2020). 10.1093/molbev/msaa089

41 Crapitto, A. J., Campbell, A., Harris, A. J. & Goldman, A. D. A consensus view of the proteome of the last universal common ancestor. Ecol Evol 12, e8930 (2022). 10.1002/ece3.8930

42 Goldman, A. D. & Kacar, B. Cofactors are Remnants of Life’s Origin and Early Evolution. J Mol Evol 89, 127–133 (2021). 10.1007/s00239-020-09988-4

43 Goldman, A. D., Beatty, J. T. & Landweber, L. F. The TIM Barrel Architecture Facilitated the Early Evolution of Protein-Mediated Metabolism. J Mol Evol 82, 17–26 (2016). 10.1007/s00239-015-9722-8

44 Rahman, K. M. T. & Butzin, N. C. Counter-on-chip for bacterial cell quantification, growth, and live-dead estimations. Sci Rep 14, 782 (2024). 10.1038/s41598-023-51014-2

45 Luef, B. et al. Diverse uncultivated ultra-small bacterial cells in groundwater. Nat Commun 6, 6372 (2015). 10.1038/ncomms7372

46 Eisen, J. A. & Hanawalt, P. C. A phylogenomic study of DNA repair genes, proteins, and processes. Mutat Res 435, 171–213 (1999). 10.1016/s0921-8777(99)00050-6

47 White, M. F. & Allers, T. DNA repair in the archaea-an emerging picture. FEMS Microbiol Rev42, 514–526 (2018). 10.1093/femsre/fuy020

48 Takeuchi, N., Hogeweg, P. & Kaneko, K. The origin of a primordial genome through spontaneous symmetry breaking. Nat Commun 8, 250 (2017). 10.1038/s41467-017-00243-x

49 Voss, N. R. & Gerstein, M. Calculation of standard atomic volumes for RNA and comparison with proteins: RNA is packed more tightly. J Mol Biol 346, 477–492 (2005). 10.1016/j.jmb.2004.11.072

50 Zamyatnin, A. A. Protein volume in solution. Prog Biophys Mol Biol 24, 107–123 (1972). 10.1016/0079-6107(72)90005-3

